## Supplementary material for "Myosin light chain 3 serves as a receptor for nervous necrosis virus entry into host cells via the macropinocytosis pathway": Fig.S1-Fig.S4;Table S1

**SUPPLEMENTAL FIGURES AND FIGURE LEGENDS**

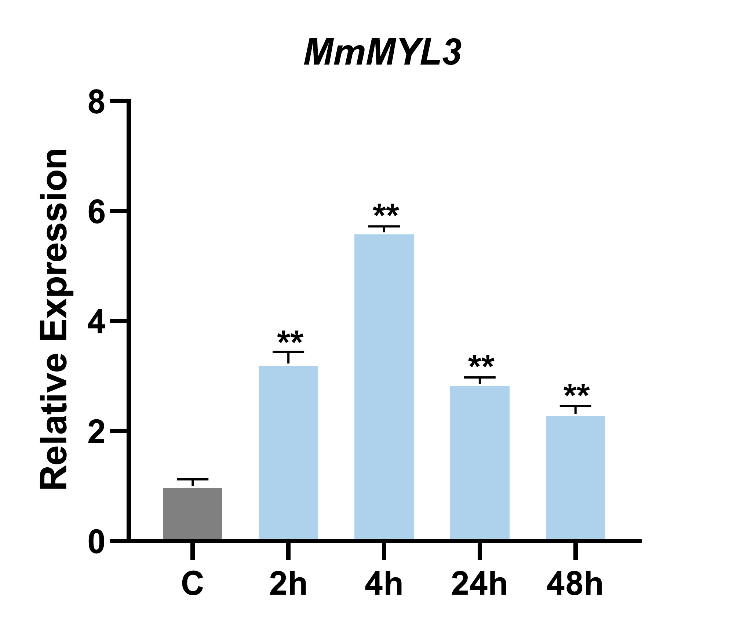

**Figure S1. Expression analysis of *MmMYL3* in hMMES1 cells at 2, 4, 24 and 48 h post RGNNV infection.** hMMES1 cells were infected with RGNNV (MOI = 1) for 2,4,24 and 48h, respectively. Then, the cells were lysed for qRT-PCR to detect the expression of *MmMYL3*. The results are presented as mean ± SD. Statistical significance was determined by an unpaired two-tailed Student’s t test. **P* < 0.05, ***P* < 0.01. Data are representative of three independent experiments.

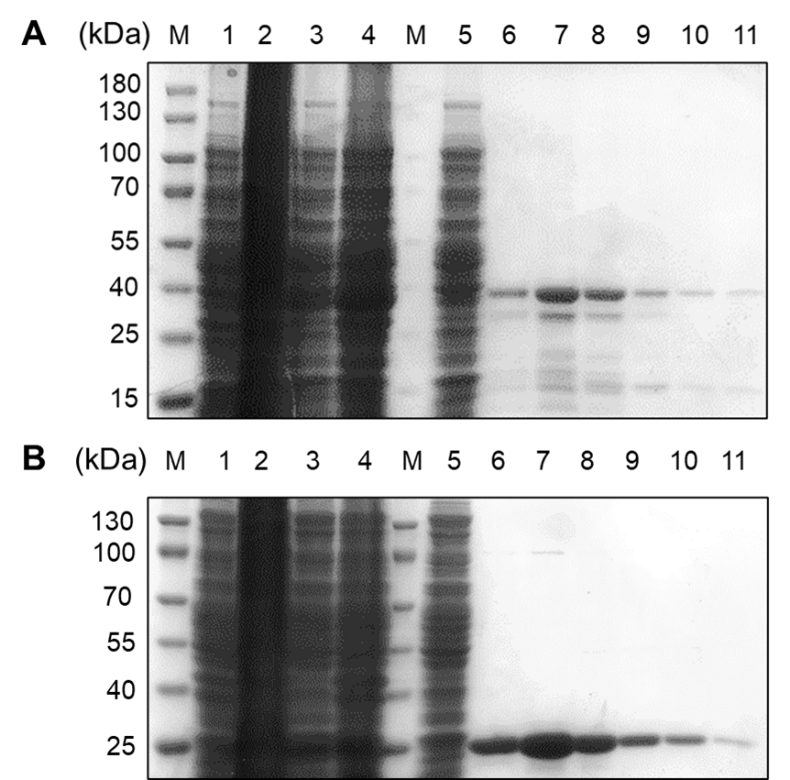

**Figure S2. Recombinant expression and purification of MmMYL3-GST (A) and GST (B).** Lane M, marker; lane 1, cell extracts before IPTG induction; lane 2, cell extracts after IPTG induction; lane 3, supernatant after centrifugation; lane 4, precipitate after centrifugation; lane 5, flow through; lane 6-11, purified recombinant.

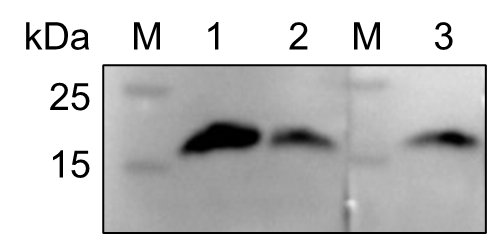

**Figure S3. MYL3 antibody validation.** Lane M, marker; lane 1, HEK 293T cell lysate; lane 2, hMMES1 cell lysate; lane 3, MmMYL3 overexpressing hMMES1 cell lysate.

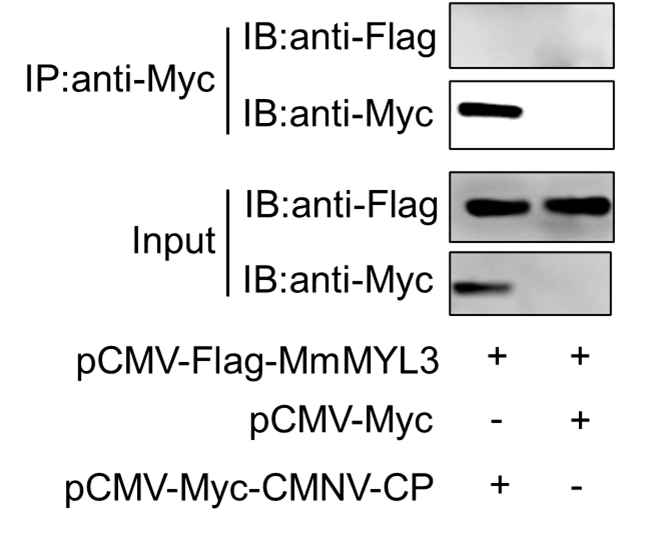

**Figure S4. MmMYL3 could not interact with CP of CMNV.** Immunoprecipitation (IP) (with anti-Myc) and immunoblot analysis (with anti-Flag and anti-Myc) of HEK293 cells transfected with MmMYL3-Flag and CMNV-CP-Myc plasmids for 48 h.

**Supplemental Table 1. The sequences of primers used in this study.**

| qRT-PCR Primers | | |
| --- | --- | --- |
| Genes | 5’ Primer | 3’ Primer |
| *MmMYL3* | ACGCCTTCCTGCCCATGCTG | CTGCCGTTCTCGTCCTCCTG |
| *MmHSP90ab1* | CCACTCCAACCGCATCTACA | TCGTCGTCAGCATCTCCTTC |
| *CP* | GTCGGCTGATACTCCTGTGTG | CTCCAGTTCCAAGGCTGTAGT |
| *RDRP* | GCTTTATGCGTGAGTGCGTC | GCTGTTTCCGTCTGTTGTGAG |
| *β-actin* | TTCAACAGCCCTGCCATGTA | CCTCCAATCCAGACAGAGTATT |
| *MmIGF1R* | AGTGCGTATCCAGACAGCTC | CGATGAGGATGGGAACGAAG |
| *MmCdc42* | CTGTAACTGTAATGATCGGCGGTGAG | CAAAGGAGGACGGCGACACG |
| *MmRac1* | GTTGTTGGAGACGGTGCTGTG | GGTAAGAAAGTGGGCGGAGT |
| *18S rRNA* | CAGCCACCCGAGATTGAGCA | TAGTAGCGACGGGCGGGTGT |
| Antireverse Primers | | |
| *CP(-)* | CGGTAGTCTCTTCAGGTGTCTC | ACAATCGTCGGCGTAGTAATC |
| Plasmid Construction Primers | | |
| pCMV-Flag/Myc-MmMYL3 | CGGAATTCCGATGACCGAGTTCACACCGGACC | CCCTCGAGTTACACAGACATGATGTGCTTG |
| pCMV-Flag/Myc-LjMYL3 | GGAATTCGGACCGAGTTCACAGCAGACCAG | GGGGTACCTTACACAGACATGATGTGCTT |
| pET-GST-MmMYL3 | CCGGAATTCATGACCGAGTTCAC | CCGCTCGAGTTACACAGACATG |
| pCMV-Myc-ΔEF-hand1 | CGGAATTCCGATGACCGAGTTCACACCGGACCAGATTGAGGGCCAGAACCCCAC | CCCTCGAGTTACACAGACATGATGTGCTTG |
| pCMV-Myc-ΔEF-hand2 | CGGAATTCCGATGACCGAGTTCACACCGG | CCCTCGAGTTACACAGACATGATGTGCTTG |
| pCMV-Flag/Myc-CP | GGAATTCGGGTACGCAAAGGTGAGAAGAAAT | ACGCGTCGACTTAGTTTCCCGAGTCAACCCTG |
| pCMV-Myc/Flag-CP-ΔARM-(34-338aa) | GGAATTCGGCGCACTGACGCACCTGTGTCTA | ACGCGTCGACTTAGTTTCCCGAGTCAACCCTG |
| pCMV-Myc/Flag-CP-Δarm-(1-33aa) | GGAATTCGGGTACGCAAAGGTGAGAAGAAAT | ATTACTACGCCGACGATTGTT |
| pCMV-Myc/Flag-CP-Δarm-(52-338aa) | AACAATCGTCGGCGTAGTAATACCAATGACGTCCATCTCTCA | ACGCGTCGACTTAGTTTCCCGAGTCAACCCTG |
| pCMV-Myc/Flag-CP-ΔLR-(1-213aa) | GGAATTCGGGTACGCAAAGGTGAGAAGAAAT | ACGCGTCGACTTGAGCGTTCCATCTCTTGAG |
| pCMV-Myc/Flag-CP-ΔLR-(221-338aa) | GTTCCATCTCTTGAGCCCATCATGACACAAGG | ACGCGTCGACTTAGTTTCCCGAGTCAACCCTG |
| pCMV-Myc/Flag-CP-ΔP-(1-220aa) | GGAATTCGGGTACGCAAAGGTGAGAAGAAAT | GGAATTCCCATCATGACACAAGGTTCC |
| pCMV-Myc/Flag-CP-ΔS-(1-51aa) | GGAATTCGGGTACGCAAAGGTGAGAAGAAAT | ACGCGTCGACTGTAACTGGATTTGGACGTGGG |
| pCMV-Myc/Flag-CP-ΔS-(214-338aa) | GATTTGGACGTGGGACACCTGAAGAGACTACCGCT | ACGCGTCGACTTAGTTTCCCGAGTCAACCCTG |
| pEGFP-MmIGF1R | CCCAAGCTTCGATGACCTGGAACGTGG | CGGGGTACCGCAGGCCGACGACTGGGG |
| pEGFP-MmIGF1R-ECD | CCCAAGCTTCGATGACCTGGAACGTGGGATC | CGGGGTACCGTAGAATGCAACGTCATCGTC |
| pEGFP-MmIGF1R-TMD | CCCAAGCTTCGATGTTGTTCATCATAATTC | CGGGGTACCGAATAAAATGGTGGTGAGGC |
| pEGFP-MmIGF1R-ICD | CCCAAGCTTCGATGTTCATCAACAAAAAGAG | CGGGGTACCGCAGGCCGACGACTGGGGC |
| pCMV-Flag/Myc-MmRac1 | CGGAATTCGGATGCAGGCCATAAAGTGTG | GGGGTACCTTAGAGGAGAGAACACTTCTTGC |
| pCMV-Flag/Myc-MmCdc42 | CGGAATTCGGATGCAGACCATCAAGTG | GGGGTACCTTATAGCAGCACACATTTGCG |
